# Pangenome analysis of SARS-CoV2 strains to Identify Potential vaccine targets by Reverse Vaccinology

**DOI:** 10.1101/2022.07.15.500170

**Authors:** Muhammad Haseeb, Afreenish Amir, Hamza Irshad

**Author notes:** Corresponding author Muhammad Haseeb Tariq, National Institute of Health, Islamabad, Pakistan.

## Abstract

**Background:** Coronavirus disease 2019 is caused by severe acute respiratory syndrome coronavirus 2 (SARS-CoV2) leads to respiratory failure and obstructive alveolar damage, which may be fatal in immunocompromised individuals. COVID-19 pandemic has severe global implications badly, and the situation in the world is depreciating with the emergence of novel variants. The aim of our study is to explore the genome of SARS-CoV2 followed by *in silico* reverse vaccinology analysis. This will help to identify the most putative vaccine candidate against the virus in a robust manner and enables cost-effective development of vaccines compared with traditional strategies.

**Methods:** The genomic sequencing data is retrieved from NCBI (Reference Sequence Number NC_045512.2). The sequences are explored through comparative genomics approaches by GENOMICS to find out the core genome. The comprehensive set of proteins obtained was employed in computational vaccinology approaches for the prediction of the best possible B and T cell epitopes through ABCpred and IEDB Analysis Resource, respectively. The multi-epitopes were further tested against human toll-like receptor and cloned in *E. coli* plasmid vector.

**Findings:** The designed Multiepitope Subunit Vaccine was non-allergenic, antigenic (0.6543), & non-toxic, with significant connections with the human leukocyte antigen (HLA) binding alleles, and collective global population coverage of 84.38%. It has 276 amino acids, consisting of an adjuvant with the aid of EAAAK linker, AAY linkers used to join the 4 CTL epitopes, GPGPG linkers used to join the 3 HTL epitopes and KK linkers used to join the 7 B-cell epitopes. MESV docking with human pathogenic toll-like receptors-3 (TLR3) exhibited a stable & high binding affinity. An in-silico codon optimization approach was used in the codon system of E. coli (strain K12) to obtain the GC-Content of *Escherichia coli* (strain K12): 50.7340272413779 and CAI-Value of the improved sequence: 0.9542834278823386. The multi-epitope vaccine’s optimized gene sequence was cloned in-silico in *E. coli* plasmid vector pET-30a (+), BamHI and HindIII restriction sites were added to the N and C-terminals of the sequence, respectively.

**Conclusion:** There is a pressing need to combat COVID-19 and we need quick and reliable approaches against Covid-19. By using In-silico approaches, we acquire an effective vaccine that could trigger adequate immune responses at the cellular and humoral level. The suggested sequences can be further validated through in vivo and in vitro experimentation.

**Statement of Significance:** Current developments in the immunological bioinformatics areas has resulted in different servers and tools that are cost and time efficient for the traditional vaccine development. Though for designing a multiple epitope vaccine the antigenic epitopes prediction of a relevant protein by immunoinformatic methods are very helpful.

## Introduction

The Wuhan city of China, became the outbreak region of origin of an unknown cause of pneumonia in December 2019 and later on, it was designated as Corona virus disease (COVID-19) by World Health Organization (WHO) in February 2020 (Zhou et al., 2020).

On basis of genetic properties, the coronavirinae family contains four genes Alpha, Beta, Gamma and Delta coronavirus. In the past twenty years the two beta coronaviruses, SARS-CoV and MERS-CoV have shown epidemic, more than 10,000 combined cases have been shown by these two (Huang et al., 2020; Wei Ji, Wei Wang, Xiaofang Zhao, Junjie Zai, n.d.). The terminology “Reverse Vaccinology” suggests a whole change of act and direction in the study of vaccines. Whole genome sequencing reformed biology including microbiology.

Especially, for the development of vaccine antigens through computer database which is very sensitive in comparison to laboratory-based assumption driven evaluation of microbes to classify elements that could display protective immunity. (Mora et al., 2003)

For the identification of vaccines, computational techniques can be employed for reverse vaccinology. For advanced experimentation, modification is crucial for their ideal use. These computational techniques help to predict antigens that are almost similar to provoke defensive responses but also describe complete detailed antigen regions and its epitopes. (Tahir ul Qamar et al., 2020)

S protein plays an important role in facilitating entry of the virus as it is most prominent and superficial protein of the coronaviruses. Previously for MERS and SARS vaccine production, the S protein, & its subunit S1 have been regularly used as vaccine antigens because of their properties to persuade neutralizing antibodies which stop infection and prevent entry into host cell. But the present coronavirus vaccines (S-protein) may have difficulties in absence of persuading full protection and likely safety issues.

Currently the vaccines of MERS/SARS were described to induce incomplete protection and neutralizing antibodies in animal models. However, it is preferred to induce sterile immunity and complete protection. Furthermore, it must be strong believed that multiple immune responses, which comprises humoral, cell-mediated, or humoral are dependable for corelates of protection than antibody titers alone. In animal models both adenovirus-based recombinant vector vaccines and whole virus vaccines expressing nucleocapsid or spike protein produced neutralizing antibody response however did not deliver full protection. Safety is major concern for us as the effectiveness and safety of the vaccination plans have not been fully verified in clinical trials of humans, so novel approaches are required to increase the safety and efficacy of COVID-19 vaccine development (Ong et al., 2020).

COVID-19 varies from earlier strains having numerous hazardous resides on Corona Virus receptor-binding region (especially Gln493) which deliver valuable communication with ACE2 human receptors. The change in closeness perhaps clarifies why this virus is more transmissible than other viruses. Main idea within all vaccinations is capability of vaccine to induce immune response in better and exponential mode than the pathogen itself. While conventional laboratory vaccines that vary on biological tests persuaded protective & neutralizing reactions in immunized animals. However, it may be hypersensitive, time consuming, costly, & foremost involving in vitro culture of infective viruses leading us to important safety issues. Thus, a cheap, antiallergic and highly efficient vaccine should be made for future. (Abdelmageed et al., 2020)

The aim of our study is to

1. Target Sars-Cov-2 by reverse vaccinology.
2. This strategy will identify the most putative vaccine candidates from many antigens and enabling cost effective development of vaccines compared with traditional strategies.

## Materials and Methodology

The overall workflow used to develop Multiepitope vaccine against SARS-CoV2 as shown in Figure *1*

**Figure 1.**
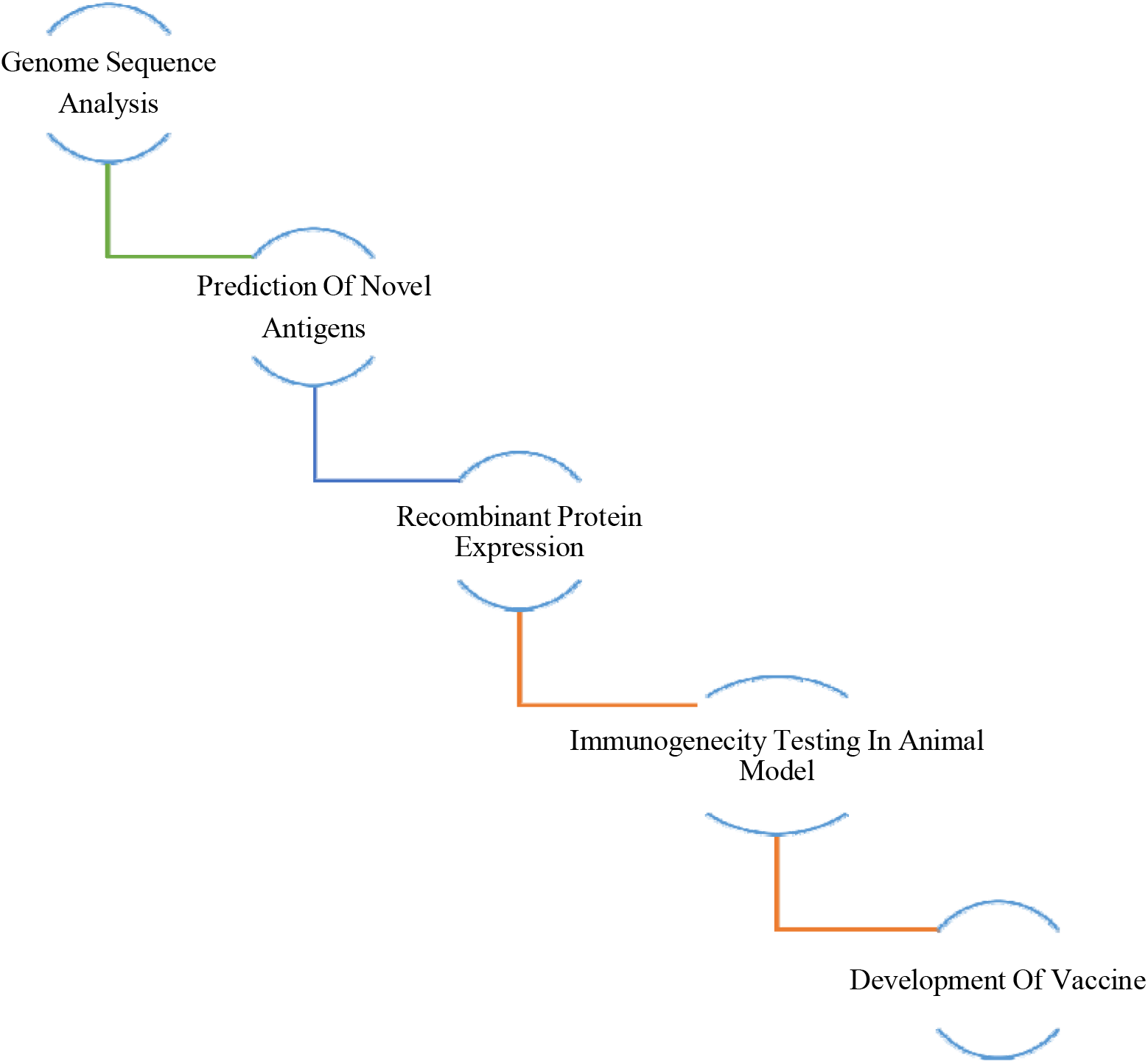
Development of the vaccines by reverse vaccinology (Kanampalliwar et al., 2013)

**Figure 2.**
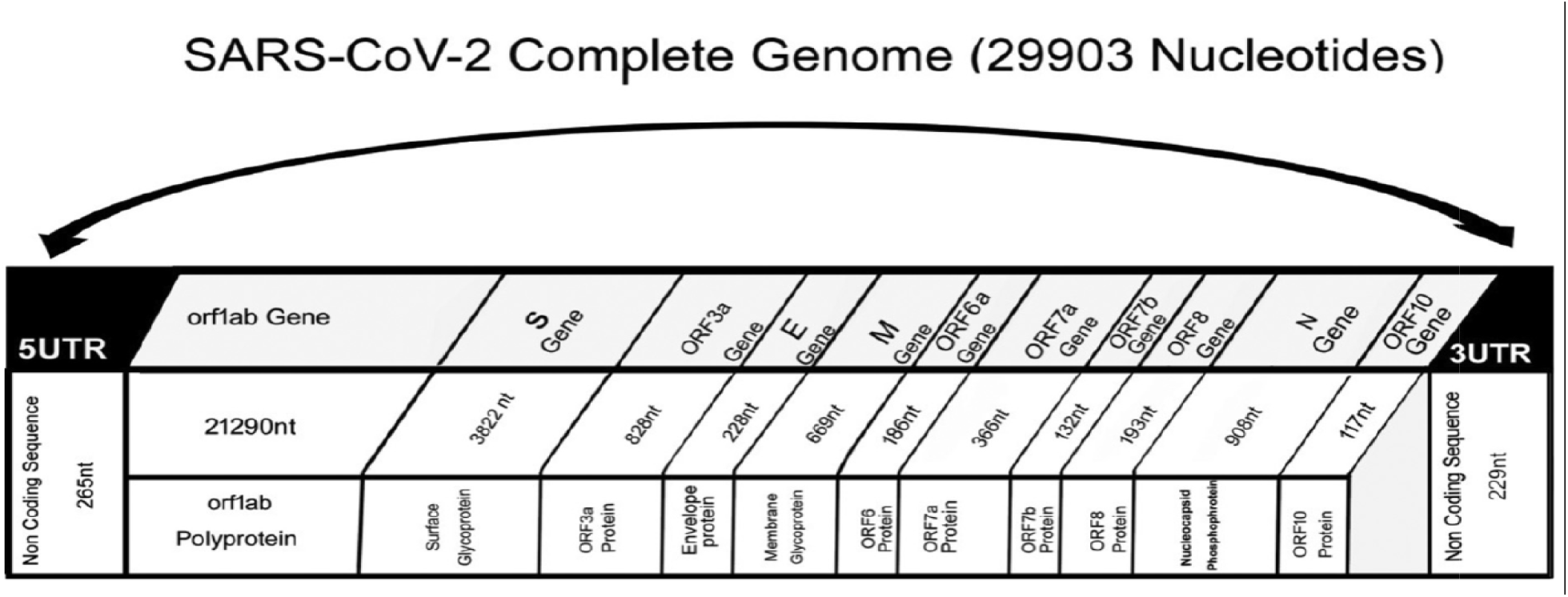
SARS-CoV-2 Complete Genome (Khailany et al., 2020)

|  |  |
| --- | --- |
| *1. Selection of Viral Strain from NCBI* |  |
| *2. Retrieval of protein sequence* |  |
| *3. Antigen protein screening and structure analysis* |  |
| **4. Identification of B-Cell Epitopes** | **5. Prediction of T-Cell Epitopes** |
| Predicting best possible B cells epitopes using IEDB tools | Trans membrane topology prediction & antigenicity analysis |
|  | Allergenicity assessment & population coverage analysis of proposed epitopes |
|  | Analysis of MHC restricted alleles |
|  | Prediction of 3D Structure of the epitopes |
|  | Molecular docking analysis within top epitopes |
| 6. Vaccine production using proposed epitopes, adjuvants, and appropriate linkers |  |
| 7. Solubility assessment, Antigenicity and Hypersensitivity of vaccine construct |  |
| 8. Physiochemical characterization and secondary structure analysis |  |
| 9. Validation, prediction, and enhancement of Tertiary structure |  |
| 10. Protein-protein docking and molecular dynamic reproduction |  |
| 11. Insilco cloning |  |

### Viral Strain selection and Retrieval of protein sequence

Firstly, proteome of SARS-CoV-2 virus were extracted from GenBank of National Center for Biotechnology Information (www.ncbi.nlm.nih.gov) and for further analysis sequences of amino acids were obtained in FASTA format. The target proteins include Surface glycoprotein, Membrane Glycoprotein, Envelop Protein, Nucleocapsid and Open Read Fragments. The chemical and physical properties of selected protein was verified by Expassy Protparam tool (https://web.expasy.org/protparam/). Only stable stability profiling proteins were further analyzed and remaining were extracted out (Gasteiger et al., 2005).

### Antigen protein screening and structure analysis

The Vaxijen 2.0 (http://www.ddg-pharmfac.net/vaxijen/VaxiJen/VaxiJen.html) software was used for protein antigenicity. For secondary structure prediction of proteins SOPMA tool (http://npsa-prabi.ibcp.fr/cgi-bin/npsa_automat.pl?page=/NPSA/npsa_sopma.html) was used. The tertiary structures details of selected proteins were predicted by using swiss (https://swissmodel.expasy.org/interactive) and Ramachandran plot (https://swift.cmbi.umcn.nl/servers/html/ramaplot.html) (Waterhouse et al., 2018) and errat results were checked through SAVES v6.0 (<u>SAVESv6.0 - Structure Validation Server (ucla.edu)</u> for protein structures verification.

### Identification of B-Cell Epitopes

B-cell epitopes which were highly antigenic and 100% conserved in all sequences of proteins were screened out and verified by ABCpred (http://crdd.osdd.net/raghava/abcpred/) (Saha & Raghava, 2006) then their Antigenicity was checked through Vixen 2 and allergenicity was checked through AllerTOP v. 2.0 (https://www.ddg-pharmfac.net/AllerTOP/). Default parameters were used (maximum distance: 6 A°; minimum score: 0.5), The conformational B-cell epitopes of the proposed MESV were estimated using the Ellipro tool (http://tools.iedb.org/ellipro/) given by IEDB-AR v.2.22. To anticipate epitopes, it relies at the residual protrusion index (PI), protein structure, and neighbor residue clustering. (Ponomarenko et al., 2008).

### Prediction of T-Cell Epitopes

IEDB Analysis Resource (http://tools.iedb.org/mhci/) was used for the prediction of T-cell HLA class I. This server predicts peptide binders to MHC molecules from protein sequences using the MHC-I binding prediction, NetMHCpan EL 4.1. (Reynisson et al., 2020). Prediction of T-cell HLA class 2 was done through IEDB Analysis Resource (http://tools.iedb.org/mhcii/). A seven-allele HLA reference set was chosen: HLA-DRB1*03:01, HLA-DRB1*07:01, HLA-DRB1*15:01, HLA-DRB3*01:01, HLA-DRB3*02:02, HLA-DRB4*01:01, HLA-DRB5*01:01. HLA-DRB1*03:01, HLA-DRB1*07:01, HLA-DRB1*15:01, HLA-DR Using the IEDB Recommended, this service predicts peptide binders to MHC molecules from protein sequences. (Wang et al., 2008). AlgPred server was used for allergenicity calculation, Non-Overlapping HTL epitopes were selected manually, Toxicity checked by ToxinPred server, Antigenicity by Vaxijen server, Interferon Gamma including ability by INF epitope server. The immunogenicity of HLA Class 1 and 2 were checked through Class I Immunogenicity (http://tools.iedb.org/immunogenicity/).

### Predict epitopes digesting enzymes

To predict epitopes digesting enzymes of epitopes Protein Digest server (http://db.systemsbiology.net:8080/proteomicsToolkit/proteinDigest.html) was used and to check toxicity of epitopes Toxin Pred (http://crdd.osdd.net/raghava/toxinpred/) was used. This tool was used to identify highly toxic or non-toxic peptides from peptides. It also predicts their toxicity along with all the important physio-chemical properties like hydrophobicity, charge pI etc. of peptides. And Toxic epitopes were excluded for further analysis (S. Gupta et al., 2013).

### Population coverage analysis of proposed epitopes

For population coverage, the Population Coverage server (http://tools.iedb.org/population/) was utilized. T lymphocytes identify a complex formed by microbe epitope and a particular major histocompatibility complex (MHC) molecule. Only individuals who have an MHC molecule able of binding that epitope will have a response to it. Denominated MHC limitation of T cell responses is the term given to this phenomenon. Human MHC (HLA) molecules are very polymorphic, with over a thousand known distinct alleles. Selecting several peptides with varying HLA binding specificities will allow peptide-based vaccinations or diagnostics to cover a larger patient community. The fact that various HLA types are expressed at drastically varying frequencies in various ethnicities further complicates the problem of population coverage in respect to MHC polymorphism. As a result, if not carefully considered, a vaccination or diagnostic might be developed with ethnically bias population coverage.

### Evaluation of Multi-epitope-based subunit vaccine (MESV)

To develop a subunit vaccine, epitopes that are (a) highly antigenic, (b) immunogenic, (c) non-allergic, (d) non-toxin, and (e) with large population coverage are typically chosen. As a result, only those epitopes were chosen for further construction of MESV using the above criteria. To enhance the immunological response, an adjuvant added to the first cytotoxic T lymphocytes (CTL) epitope using the EAAAK linker. After confirming their interaction compatibility, further epitopes were connected using AAY, GPGPG, and KK linkers to retain their separate immunogenic activity. Because it is a 45 amino-acid-long peptide that works as both an immunomodulation and as well as antimicrobial agent, β-defensin was adopted as an adjuvant in this study (Hoover et al., 2003).

First, using default parameters, Blastp analysis was used to check that predicted MESV sequence is not similar to the proteome of Homo sapiens.(Mahram & Herbordt, 2010) Non-homologous proteins are those that have less than 37% homology. The Protparam tool was used to access the physiochemical characteristics of the designed MESV (Gasteiger et al., 2005a). Depending on the amino acid assumptions used in the pk, Protparam estimates several physiochemical characteristics such as half-life, theoretical isoelectric point [pI], instability index, grand average hydropathy, and aliphatic index (Bjellqvist et al., 1993). The AllerTOP v.2.0 server was used to determine the allergenicity of the MESV construct. The secondary structure of the MESV construct was examined by using the PSIPRED workbench. (Mahram & Herbordt, 2010). The alpha helices, extended chain, degree of beta folds, and random coil features of the vaccine were also analyzed in this test.

Because the proposed MESV comprised a sequence of epitopes and no relevant template was available, MESV’s 3D structure was anticipated using the CABS fold server’s de novo modelling approach (http://biocomp.chem.uw.edu.pl/CABSfold/) (Blaszczyk et al., 2013). The CABS modelling technique, which merges with a multi-scale modelling pipeline with exchange replica Monte Carlo scheme, is used to build this server. A galaxy refine server was used to modify the predicted MESV 3D structure (Heo et al., 2013). The RAMPAGE server (http://mordred.bioc.cam.ac.uk/rapper/rampage.php) was used to analyze the Ramachandran plot http://mordred.bioc.cam.ac.uk/~rapper/rampage.php (Lovell et al., 2003), The PROSA web server was to check the refinement of the MESV structure, followed by analysis of structural validation (Wiederstein & Sippl, 2007b). The MESV structure’s prediction of unbounded interactions was also evaluated using the ERRAT server (https://servicesn.mbi.ucla.edu/ERRAT/) (Qamar et al., 2020).

### Molecular docking of MESV with human immune receptors

The interaction between the antigenic molecule and the immune receptor molecule is critical for the proper evocation of immune response. The interaction between the MESV construct and human immune receptors was evaluated using molecular docking. TLR3 (Toll-like Receptors-3) has been studied extensively, and research have revealed that it plays a critical role in the generation of antiviral immune responses. The MESV docking with TLR3 was conducted using GRAMM-X (http://vakser.compbio.ku.edu/resources/gramm/grammx/) (PDB ID: 1ZIW) for visualization pymol was used to see the docked complexes (DeLano, 2002). Furthermore, a web server called PDBsum (http://www.ebi.ac.uk/thornton-srv/databases/cgi-bin/pdbsum/GetPage.pl?pdbcode=index.html) was used to construct a typical sketch of interactions among docked proteins. It analyses docked molecules’ among protein-protein interactions. (Laskowski et al., 2018).

### Immunogenicity evaluation of the vaccine construct

To validate the immunological responses of the constructed MESV, an in silico immune simulation was performed using the C-ImmSim 10.1 server (http://150.146.2.1/C-IMMSIM/index.php?page=0). The three primary components of the functional mammalian system (thymus, lymph node, and bone marrow) are all simulated in C-ImmSim (Rapin et al., 2010b). The input parameters for the immune simulations are as follows: volume (10), HLA (A0101, A0101, B0702, B0702, DRB1_0101, DRB1_0101), random seed (12345), number of steps (100), number of injections set to 1. The remaining settings were presumed to be at default.

### In silico cloning and codon optimization

If the use of codon is different in both organisms, codon optimization is a way to enhance the translation efficacy of foreign genes in the host. After a comprehensive review of MESV features and immunological response, codon optimization and in silico cloning were carried out. The java codon adaptation tool (http://www.jcat.de/) (Grote et al., 2005) was used for MESV codon optimization to make this tool compatible with the extensively used prokaryotic expression system, E. coli K12 (Smith et al., 1987). The other options were chosen to avoid: (i) rho-independent transcription termination, (ii) prokaryote ribosome binding sites, and (iii) restriction enzyme cleavage sites. The GC (guanine and cytosine) contents, as well as the codon adaptation index (CAI) (Sharp & Li, 1987), were analyzed. Sticky ends of HindIII and BamHI restriction sites were attached to the start/N terminal and end/C terminal of the modified MESV sequence, respectively, to enable restriction and cloning. To assure in vitro expression, the modified nucleotide sequence of MESV was also cloned into the E. coli pET30a (+) vector using the SnapGene tool (https://www.snapgene.com/).

## Results

Firstly, proteome of SARS-CoV-2 virus was taken from GenBank of NCBI (www.ncbi.nlm.nih.gov) and for further analysis sequences of amino acids were obtained in FASTA format. The target proteins include Surface glycoprotein, Membrane Glycoprotein, Envelop Protein, Nucleocapsid. The chemical and physical properties of selected protein was verified by Expassy Protparam tool (https://web.expasy.org/protparam/) as shown in Table *1*. Only stable stability profiling proteins were further analyzed and remaining were extracted out (Gasteiger et al., 2005b).

**Table 1.** Physiochemical properties of the SARS-COV-2 proteins.

| Proteins | Molecular Weight | Theoretical pI | Instability index | Stability Profiling | Aliphatic Index |
| --- | --- | --- | --- | --- | --- |
| Surface | 141120.43 | 6.32 | 32.86 | Stable | 84.67 |
| Membrane | 25146.62 | 9.51 | 39.14 | Stable | 120.86 |
| Envelop | 8365.04 | 8.57 | 36.68 | Stable | 144.00 |
| Nucleocapsid | 45696.82 | 10.09 | 55.81 | Unstable | 52.53 |
| ORF1A | 490043.01 | 6.02 | 35.02 | Stable | 89.16 |
| ORF1AB | 794127.93 | 6.31 | 33.35 | Stable | 87.03 |
| ORF3A | 31122.94 | 5.55 | 32.96 | Stable | 103.42 |
| ORF6 | 7272.54 | 4.60 | 31.16 | Stable | 130.98 |
| ORF7A | 13744.17 | 8.23 | 48.66 | Unstable | 100.74 |
| ORF7B | 5180.27 | 4.17 | 50.96 | Unstable | 156.51 |
| ORF8 | 13831.01 | 5.42 | 45.79 | Unstable | 97.36 |
| ORF10 | 4449.23 | 7.93 | 16.06 | Stable | 107.63 |

The Vaxijen 2.0 (http://www.ddg-pharmfac.net/vaxijen/VaxiJen/VaxiJen.html) software was used for protein antigenicity, Surface protein, ORF1A, ORF1AB, ORF3A proteins were excluded due to low antigenic values less than 0.5, as shown in Table *2*.

**Table 2.** Antigenic prediction of SARS-COV-2 proteins.

| Proteins | Antigen Prediction |
| --- | --- |
| Surface | 0.463 |
| Membrane | 0.510 |
| Envelop | 0.6025 |
| ORF1A | 0.4794 |
| ORF1AB | 0.4625 |
| ORF3A | 0.4945 |
| ORF6 | 0.6131 |
| ORF10 | 0.7185 |

For secondary structure prediction of proteins SOPMA tool (http://npsa-prabi.ibcp.fr/cgi-bin/npsa_automat.pl?page=/NPSA/npsa_sopma.html) was used as shown in Table *3***Error! Reference source not found**.. (Blanchet et al., 2000)

**Table 3.** Secondary structure of COV-2 proteins.

| Proteins | Sequence Length | Alpha Helix | Beta Turn | Random Coil | Extended Strand |
| --- | --- | --- | --- | --- | --- |
| Membrane | 222 | 34.86% | 6.76% | 37.39% | 21.17% |
| Envelop | 75 | 45.3% | 9.33% | 20.0% | 26.67% |
| ORF6 | 61 | 70.49% | 8.20% | 11.48% | 9.84% |
| ORF10 | 38 | 28.95% | 5.26% | 28.95% | 36.84% |

The tertiary structures details of selected proteins were predicted by using swiss (https://swissmodel.expasy.org/interactive) and Ramachandran plot (https://swift.cmbi.umcn.nl/servers/html/ramaplot.html) as shown in Table *4*. (Waterhouse et al., 2018) and errat results were checked through SAVES v6.0 (<u>SAVESv6.0 - Structure Validation Server (ucla.edu)</u>). ORF10 was excluded due to non-homologous structure and no such template was found.

**Table 4.**
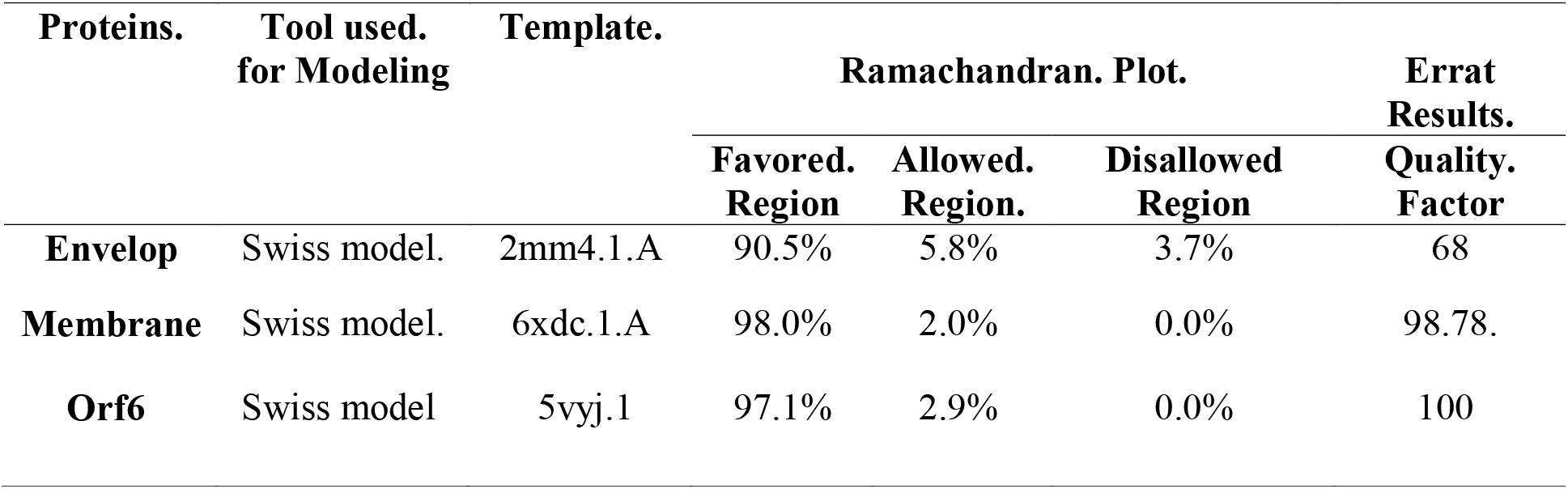
3D structural. Details of COV-2 proteins.

B-cell epitopes which are screened out are highly antigenic and 100% conserved in all sequences of proteins were verified by ABCpred (http://crdd.osdd.net/raghava/abcpred/) then their Antigenicity was checked through Vixen 2 and allergenicity was checked through AllerTOP v. 2.0 (https://www.ddg-pharmfac.net/AllerTOP/) those epitopes were not selected further which were probably non-antigen and allergen and non-overlapping. Furthermore total 8 conformational linear epitopes of all the proteins are in shown in Table 5

**Table 5:**
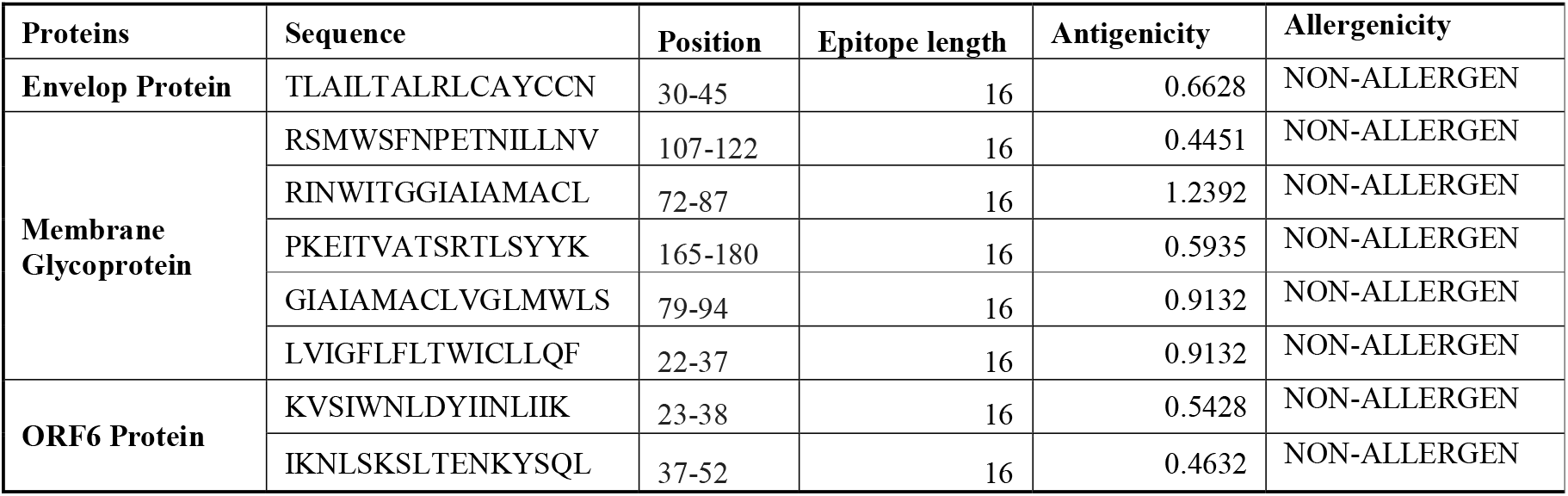
Predicted epitopes of B-cell of Envelop, Membrane & ORF6 proteins.

### T-Cell Epitopes

Prediction of T-cell HLA class I was done through IEDB Analysis Resource (http://tools.iedb.org/mhci/). This server predicts peptide binders to MHC molecules from protein sequences using the MHC-I binding prediction, NetMHCpan EL 4.1. We have selected all HLA class I alleles from the selection panel of IEDB Analysis Resource for prediction of epitopes of HLA class I. The selected epitopes were Immunogenic, Antigenic and Non-toxic (Reynisson et al., 2020).

Prediction of T-cell HLA class 2 was done through IEDB Analysis Resource (http://tools.iedb.org/mhcii/). We had selected 7-allele HLA reference set: HLA-DRB1*03:01, HLA-DRB1*07:01, HLA-DRB1*15:01, HLA-DRB3*01:01, HLA-DRB3*02:02, HLA-DRB4*01:01, HLA-DRB5*01:01. The one with low adjusted rank were good binders. This server predicts peptide binders to MHC molecules from protein sequences using the IEDB Recommended (Wang et al., 2008). The selected epitopes were non-allergic, non-overlapping, non-toxic, Antigenic. AlgPred server was used for allergenicity calculation, Non-Overlapping HTL epitopes were selected manually, Toxicity checked by ToxinPred server, Antigenicity by Vaxijen server, Interferon Gamma including ability by INF epitope server. The immunogenicity of HLA Class 1 and 2 were checked through Class I Immunogenicity (http://tools.iedb.org/immunogenicity/).

### Predict epitopes digesting enzymes

To predict epitopes digesting enzymes of epitopes Protein Digest server (http://db.systemsbiology.net:8080/proteomicsToolkit/proteinDigest.html) was used and to check toxicity of epitopes Toxin Pred (http://crdd.osdd.net/raghava/toxinpred/). This tool was used to identify highly toxic or non-toxic peptides from peptides. It also predicts their toxicity along with all the important physio-chemical properties like hydrophobicity, charge pI etc. of peptides. And toxic epitopes were excluded for further analysis (Gupta et al., 2013).

### Population Coverage

Population Coverage server (http://tools.iedb.org/population/) was used for population coverage. T lymphocytes identify a complex formed by a pathogen-derived epitope and a specific major histocompatibility complex (MHC) molecule. Only individuals who have an MHC molecule capable of binding that epitope will have a response to it. Denominated MHC limitation of T cell responses is the term given to this phenomenon. Human MHC (HLA) molecules are very polymorphic, with over a thousand known distinct alleles. Selecting several peptides with varying HLA binding specificities will allow peptide-based vaccinations or diagnostics to cover a larger patient community. The fact that various HLA types are expressed at drastically varying frequencies in different ethnicities further complicates the problem of population coverage in respect to MHC polymorphism. As a result, if not carefully considered, a vaccination or diagnostic might be developed with ethnically bias population coverage as shown in Table 6

**Table 6.** Population coverage of selected epitopes of Class I, Class II and Class combined

| population/area | Class I |  |  | Class II |  |  | Class combined |  |  |
| --- | --- | --- | --- | --- | --- | --- | --- | --- | --- |
|  | coverage | average_hitb | pc90c | coverage | average_hitb | pc90c | coverage | average_hitb | pc90c |
| China | 41.21% | 0.51 | 0.17 | 33.37% | 1.22 | 0.15 | 60.83% | 1.73 | 0.26 |
| East Asia | 50.68% | 0.72 | 0.2 | 38.36% | 1.44 | 0.16 | 69.60% | 2.16 | 0.33 |
| Europe | 58.59% | 0.75 | 0.24 | 74.58% | 3.12 | 0.39 | 89.48% | 3.87 | 0.95 |
| France | 54.73% | 0.71 | 0.22 | 77.02% | 3.07 | 0.44 | 89.60% | 3.78 | 0.96 |
| Germany | 61.05% | 0.78 | 0.26 | 83.63% | 3.63 | 0.61 | 93.62% | 4.41 | 1.34 |
| India | 38.70% | 0.62 | 0.16 | 65.14% | 2.78 | 0.29 | 78.63% | 3.4 | 0.47 |
| Iran | 56.57% | 0.74 | 0.23 | 54.51% | 2.3 | 0.22 | 80.24% | 3.04 | 0.51 |
| Italy | 49.92% | 0.59 | 0.2 | 20.93% | 0.77 | 0.13 | 60.40% | 1.36 | 0.25 |
| Japan | 52.89% | 0.8 | 0.21 | 26.97% | 0.96 | 0.14 | 65.60% | 1.77 | 0.29 |
| Northeast Asia | 40.68% | 0.5 | 0.17 | 33.37% | 1.22 | 0.15 | 60.47% | 1.72 | 0.25 |
| Pakistan | 51.90% | 0.79 | 0.21 | 2.19% | 0.07 | 0.31 | 52.96% | 0.85 | 0.21 |
| South Asia | 45.72% | 0.71 | 0.18 | 65.61% | 2.8 | 0.29 | 81.33% | 3.52 | 0.54 |
| Spain | 55.20% | 0.67 | 0.22 | 69.56% | 2.97 | 0.33 | 86.36% | 3.64 | 0.73 |
| Turkey | 44.80% | 0.45 | 0.18 | 59.80% | 2.49 | 0.25 | 77.81% | 2.94 | 0.45 |
| Zambia | 80.49% | 1.09 | 0.51 | 0.00% | 0 | 0 | 80.49% | 1.09 | 0.51 |
| Zimbabwe | 56.70% | 0.67 | 0.23 | 50.58% | 2.09 | 0.61 | 78.60% | 2.76 | 0.47 |
| World | 55.34% | 0.72 | 0.22 | 65.03% | 2.54 | 0.29 | 84.38% | 3.26 | 0.64 |
| Average | 49.76 | 0.65 | 0.21 | 44.08 | 1.79 | 0.25 | 71.31 | 2.44 | 0.49 |
| Standard deviation | 11.99 | 0.18 | 0.08 | 25.59 | 1.1 | 0.16 | 17.14 | 1.14 | 0.3 |

### Selection of MESV

Only those epitopes were chosen for further construction of MESV which are a) highly antigenic, (b) immunogenic, (c) non-allergenic, (d) non-toxic, and (e) with large population coverage are typically chosen. To enhance the immunological response, an adjuvant was added to the first cytotoxic T lymphocytes (CTL) epitope using the EAAAK linker. After confirming their interaction compatibility, further epitopes were connected using AAY, GPGPG, and KK linkers to retain their separate immunogenic activity (Hoover et al., 2003) as shown in Figure *3*

**Figure 3.**
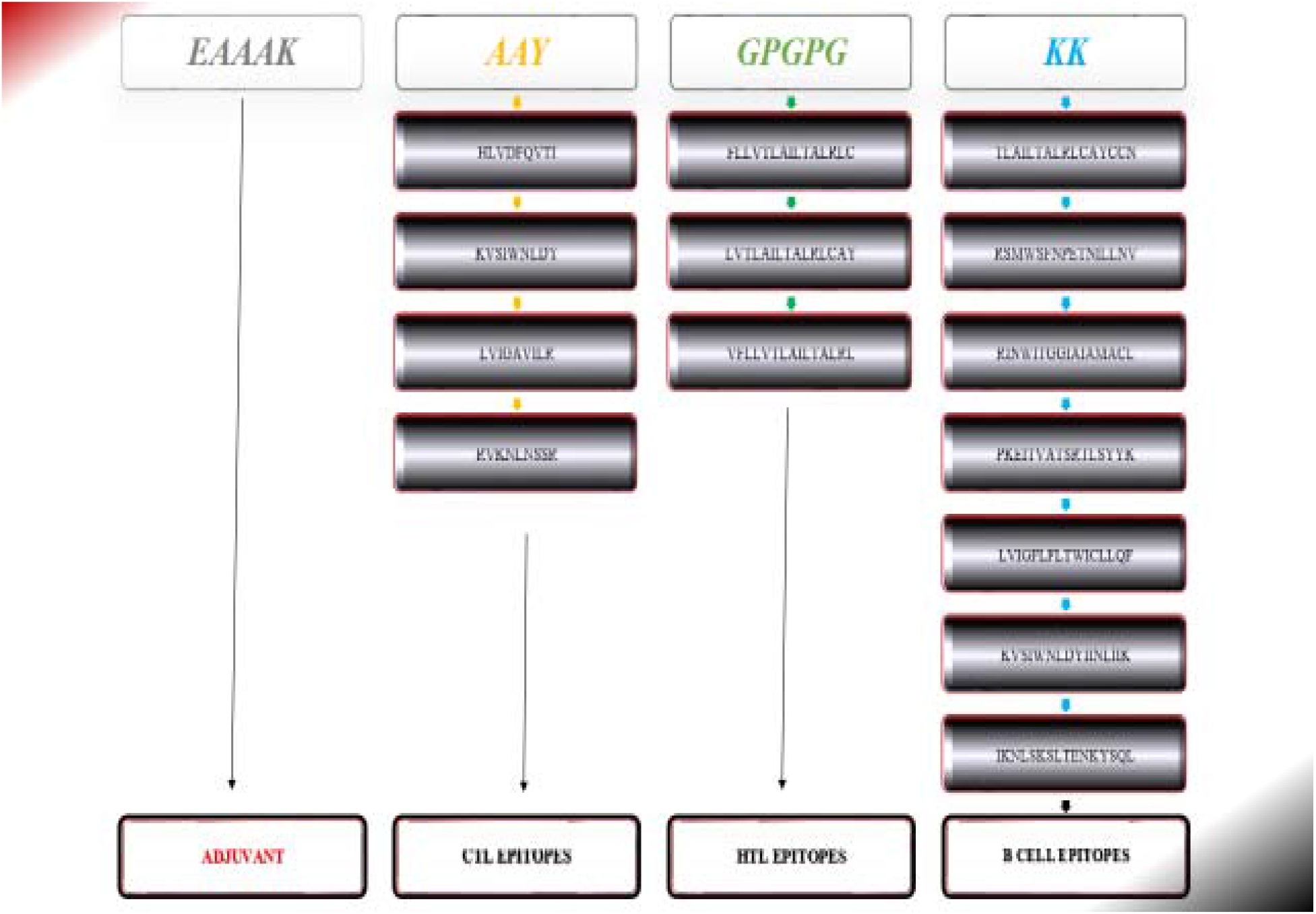
Graphic representation of MESV is a 276-amino-acid construct that includes an adjuvant (orange) attached to the N-terminal of MEV through the EAAAK linker (yellow). CTL epitopes were joined with AAY linkers (blue), HTL epitopes with GPGPG linkers (green), and B-cell epitopes with KK linkers (grey).

### Analysis of MESV

The designed sequence was then verified for Allergenicity, Toxicity, and Antigenicity. The designed MESV was highly antigenic (0.6543), non-allergenic, and non-toxic. The molecular weight of designed vaccine was Molecular weight: 31204.78 kDa and Theoretical pI: 10.13. The estimated half-life was hours (mammalian reticulocytes, in vitro), >20 hours (yeast, in vivo) >10 hours (Escherichia coli, in vivo). The instability index (II) is computed to be 33.90, aliphatic index was 120.21, and Grand average of hydropathicity (GRAVY) was 0.329. This classifies the protein as stable. The secondary structure shows that Alpha helix is 41.79%, Beta turn is 7.50% and Extended Strand is 28.21%. ProSA-web (https://prosa.services.came.sbg.ac.at/prosa.php) was used for Z-Score which was -0.37, 3D structure of the protein and Overall quality of model was checked. (Virology et al., 2017; Wiederstein & Sippl, 2007a) as shown in Figure 4

**Figure 4.**
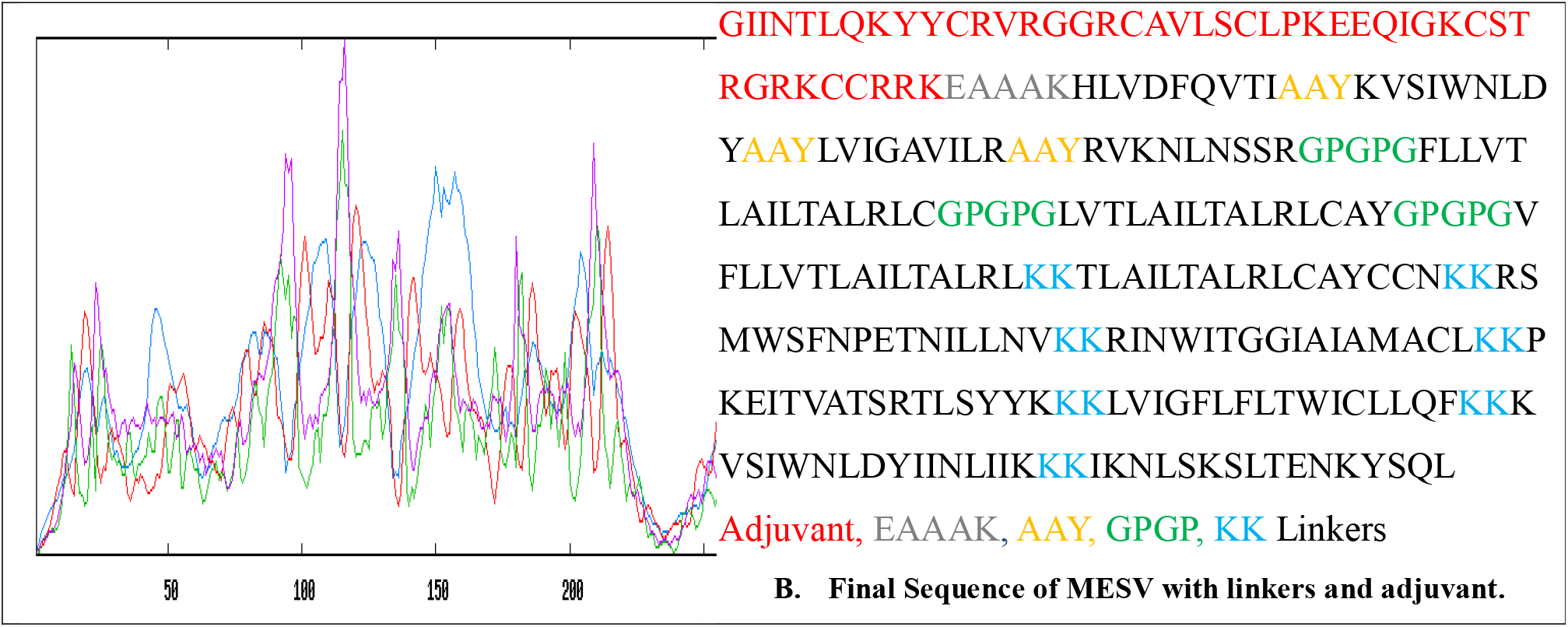
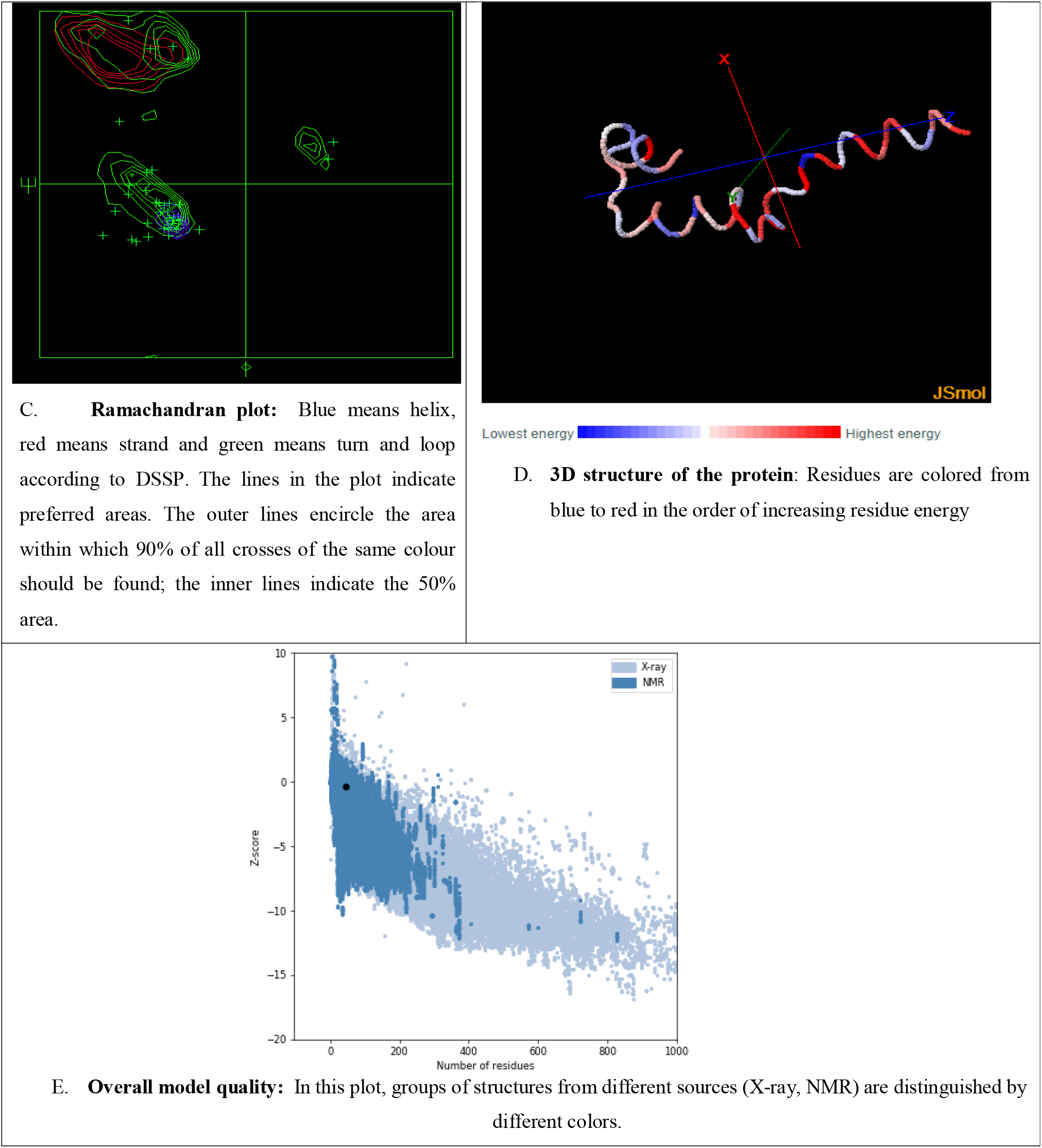
A Secondary Structure of MESV B. Final Sequence of MESV C. Ramachandran plot D. 3D structure of the protein E. Overall model quality

### Molecular docking of MESV with human immune receptors

Molecular Docking was done to evaluate the interface between human immune receptors and MESV Construct. The interaction between the antigenic molecule & immune receptor molecule is extremely important. TLR3 (Toll-like Receptors-3) has been studied extensively, & have revealed that it plays a critical role in the development of antiviral immune responses. The MESV docking with TLR3 was done by using GRAMM-X (http://vakser.compbio.ku.edu/resources/gramm/grammx/) (PDB ID: 1ZIW). Visualization of the docked complexes was done by using Pymol. Furthermore, a web server called PDBsum (http://www.ebi.ac.uk/thornton-srv/databases/cgibin/pdbsum/GetPage.pl?pdbcode=index.html) was used to generate the traditional schematic of interactions between docked proteins. It analyzes the protein-protein interactions among docked molecules. Dockthor https://dockthor.lncc.br/v2/index.php?tab=DOCKING&page=RESULTS&jobId=MolecularXdocking_60426ddd594d1 (Guedes et al., 2021; Lohning et al., 2017) as shown in Figure *5*.

**Figure 5.**
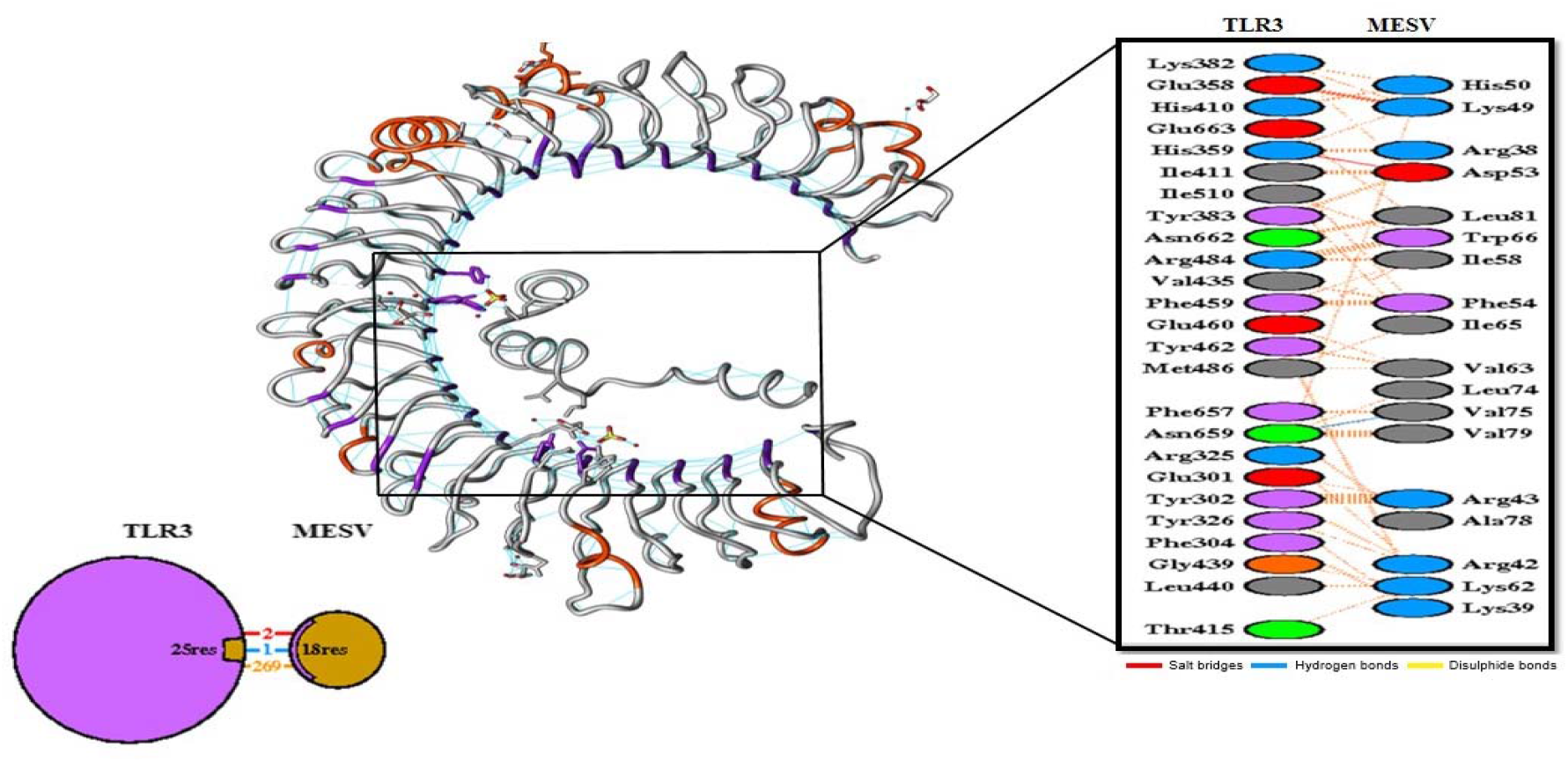
the number of H-bond lines between any two residues indicates the number of potential hydrogen bonds between them. For non-bonded contacts, which can be plentiful, the width of the striped line is proportional to the number of atomic contacts.

### Molecular dynamics simulation

The vaccine construct with the best molecular docking study findings gone through a molecular dynamics simulation study. For the molecular dynamics simulation analysis, the iMODS web-server (http://imods.Chaconlab.org/) was utilized, which is a fast, free molecular dynamics simulation server for specifying and quantifying protein flexibility as shown in Figure 6.

**Figure 6.**
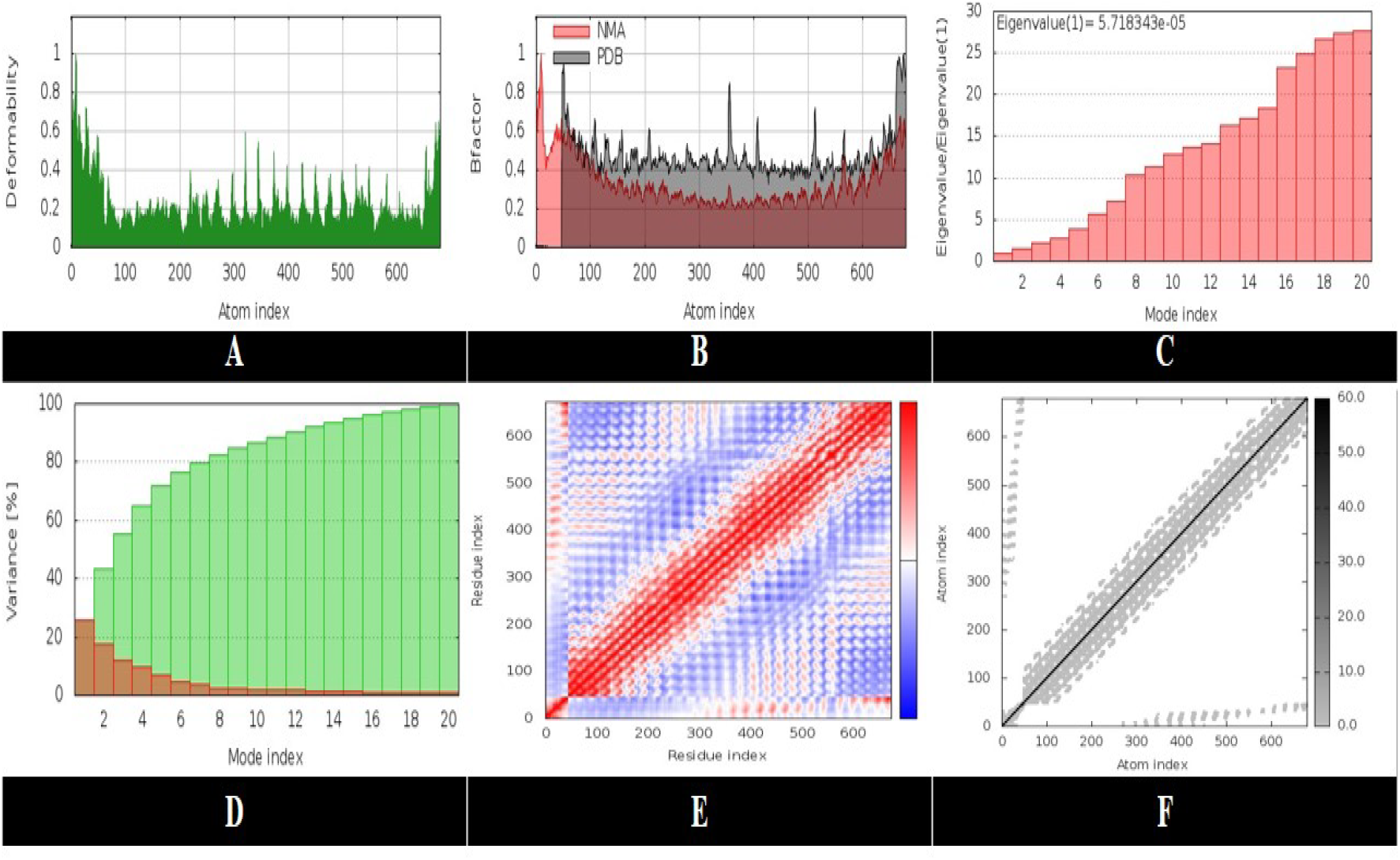
Molecular dynamics simulation of constructed MESV.

### Immune simulation

The vaccine formulation immune response profile was reported using the in silico technique C-ImmSim, through an online simulation server (http://150.146.2.1/C-IMMSIM/index.php). The humoral and cellular responses of a mammalian immune system to a vaccine construct are defined by C-ImmSim. Three doses of the preventive Covid vaccine’s target product profile were given at various intervals of four weeks. With time periods of 1, 84, and 170, all simulation settings were set to default. The simulation volume and steps were both set to ten and one thousand, respectively (random seed = 12345 with LPS-free vaccination injection)(Rapin et al., 2010a) as shown in Figure 7.

**Figure 7.**
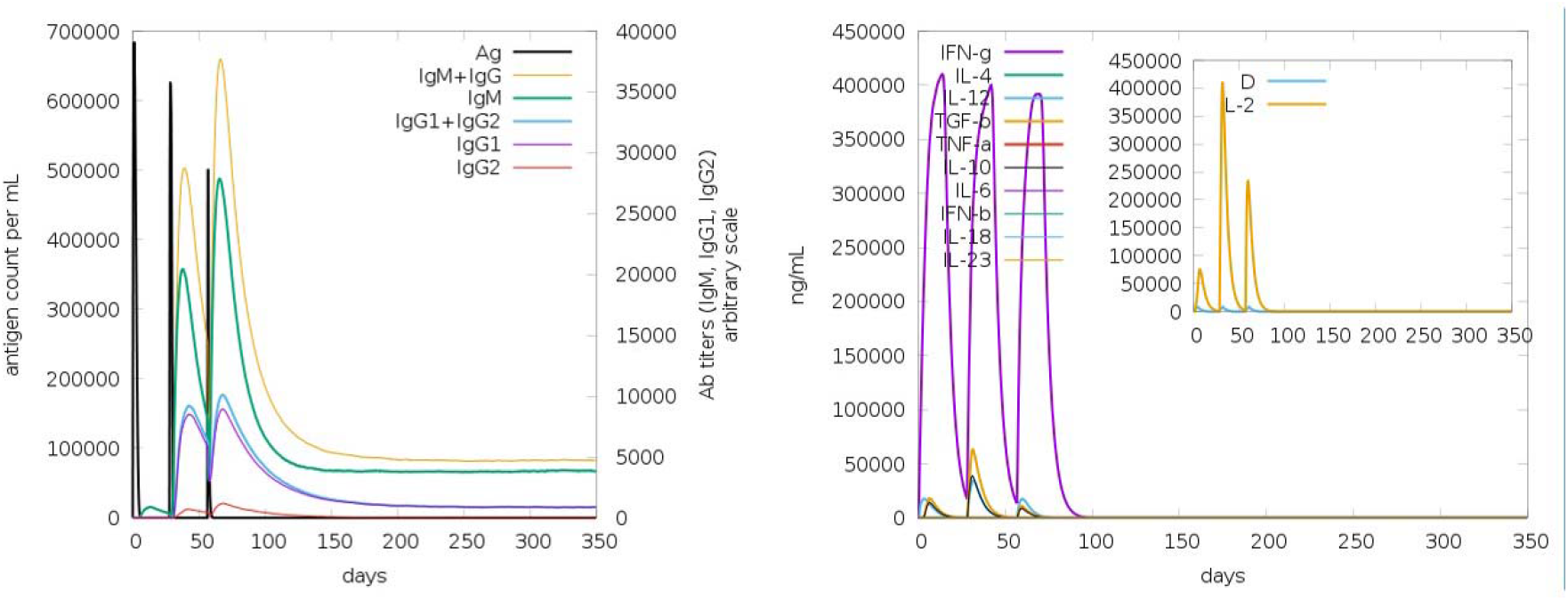
Immune stimulation of constructed MESV

### In-Silico Cloning & Codon Enhancement

To enhance recombinant protein expression, a codon optimization approach was applied. Because of the genetic code’s degeneracy, most amino acids may be encoded by several codons, which needs codon optimization. In the codon system of E. coli (strain K12), the Java Codon Adaptation Tool (JCat) server (http://www.prodoric.de/JCat) was used to get the codon adaptation index (CAI) values and GC contents to evaluate the levels of protein synthesis. The GC-Content of Escherichia coli (strain K12): 50.7340272413779 and CAI-Value of the improved sequence: 0.9542834278823386. The optimum CAI value is 1.0, while a score of > 0.95 is considered good, and the GC content ranges from 30 to 70%. Beyond this range, there are negative impacts on translation and transcriptional efficiency. The optimized gene sequence for the multi-epitope vaccine was cloned in the E. coli plasmid vector pET-30a (+), with BamHI and HindIII restriction sites added to the N and C terminals, respectively. Finally, using the Snap Gene programme (https://www.snapgene.com/free-trial/), the optimized sequence of the final vaccine construct (containing restriction sites) was introduced into the plasmid vector pET-30a (+) to validate the vaccine’s expression as shown in Figure 8.

**Figure 8.**
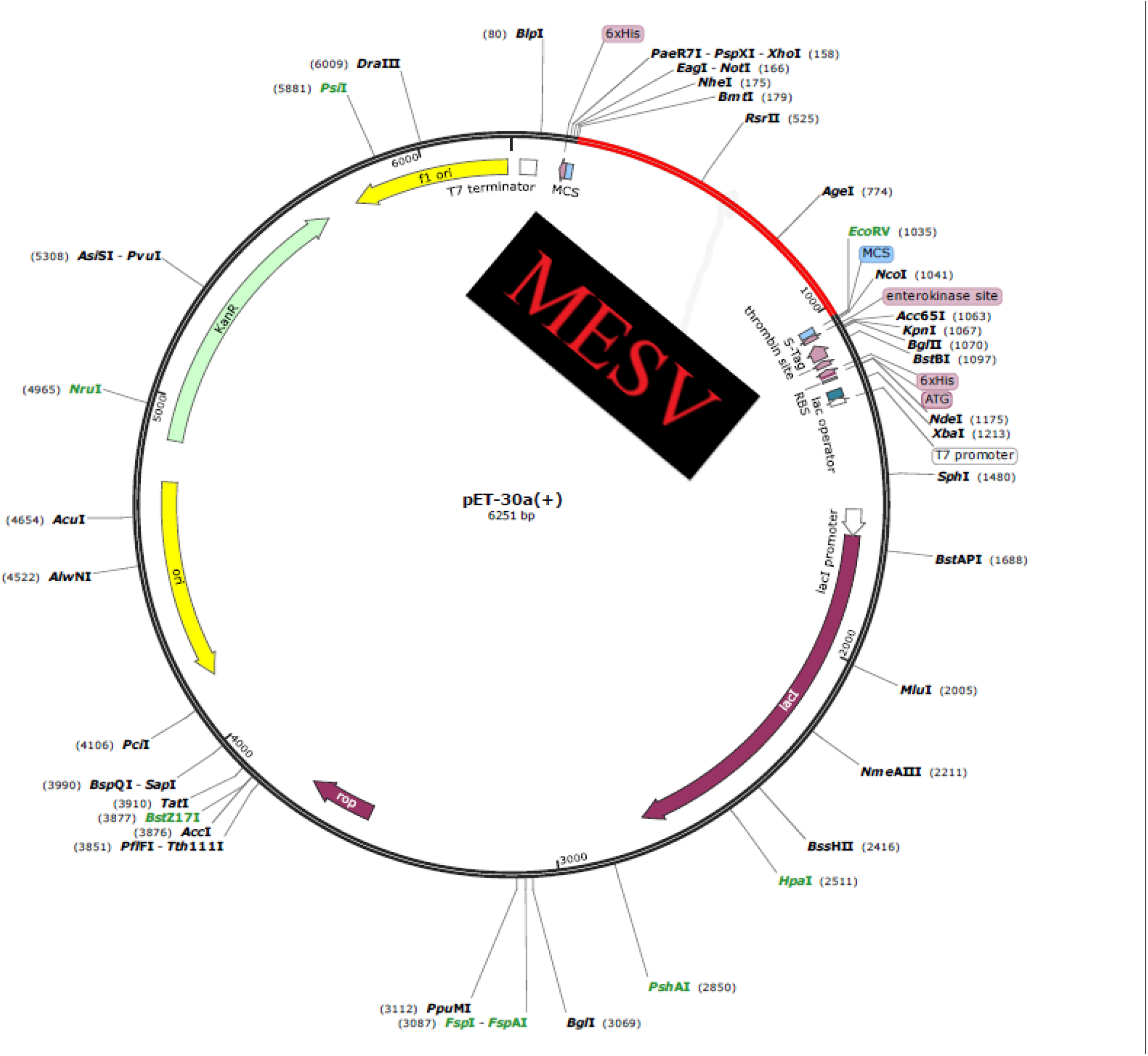
In-Silico Cloning & Codon Enhancement of constructed MESV into plasmid

## Discussion

Corona virus have long been disregarded as inconsequential microorganisms that cause human “colds.” Two particularly pathogenic of corona virus, SARS-CoV and MERS-CoV, arose from cattle reservoirs in the twenty-first century and caused devastating epidemics. Recently, a new strain of CoV, formally known as SARS-COV-2, was discovered, triggering a fatal global epidemic of COVID-19 due to the constantly developing mutations, the pandemic’s final dimensions and impact are presently unknown (Abramo et al., 2012). The new virus rapidly infects host cells after recombination of several virus genome particles. There is yet no effective treatment for the infection. Infections with COVID-19 is a major cause of illness and mortality around the world. Unfortunately, the lack of COVID-19 vaccines has resulted in the loss of countless valuable lives in various parts of the world. COVID-19’s advent has resulted in a major global medical burden, necessitating immediate prevention actions. Researchers have been collecting data on Coronavirus in order in order to better recognize the illness’s transmission, pathophysiology, and biology in order to eradicate the disease (Douglas et al., 2020). The rapid growth of databases of structural and genomic information, in combination with computational methods, contributes to the development and discovery of novel vaccination candidates. Recent advancements in bioinformatics has resulted in a variety of tools and web servers that can aid in the development of conventional vaccinations by reducing the time and cost. The creation of successful Multiepitope vaccines is difficult due to difficulties in selecting good antigen candidates and immunodominant epitopes. As a result, immunoinformatics techniques for predicting suitable antigenic epitopes of a targeted protein are critical for designing a MESV (Nain et al., 2020).

The development of epitope-based vaccines targeting the structural proteins (M and E) of the SARS-CoV-2 was investigated in this study. These proteins are essential for the replication cycle and the structure of virus particles (Kirchdoerfer et al., 2016; Siu et al., 2008; Song et al., 2004). M and E proteins are required for viral entrance, replication, and particle assembly within human cells (Alsaadi & Jones, 2019; Schoeman et al., 2020). The target proteins’ T- and B-cell epitopes were expected to aid the host’s immunological response. The research examined at proteins at the primary, secondary, and tertiary structural levels. ABCPred predicted B-cell conserved epitopes using the IEDB analysis database. Pymol was used to visualize the position of epitopes on 3D protein structures. To determine discontinuous epitopes, the DiscoTop server was utilized. Allergenicity, toxicity, and physiochemical properties of anticipated epitopes were evaluated to increase specificity and selectivity. The peptides predicted throughout the research were stable and safe to use, according to digestion analysis.

A suitable MESV should have B-cell, HTL, and CTL epitopes and induce efficient antiviral responses against a specific virus (Zhang, 2018). Subunit vaccines of SARS-CoV-2 were created by a few groups, however they only use single protein in vaccine formulation (Abdelmageed et al., 2020; Abraham Peele et al., 2021; Bhattacharya et al., 2020) and the exclusive usage of CTL epitopes without regard for the significance of B cell & HTL epitopes (Mishra, 2020). We have included B-cell epitopes from numerous structural proteins in addition T-cell epitopes are also important because they play a role in antibody formation and influencing its efficacy (Cooper & Nemerow, 1984). Furthermore, antigens can quickly overcome the humoral response of memory B-cells, whereas cell-mediated immunity (T-cell immunity) often leads to long-term immunity (Bacchetta et al., 2005). CTL prevents pathogen spread by secreting specific antiviral cytokines and identifying and destroying infected cells (Pohl et al., 2013). As a result, the current vaccine design exceeds previously reported constructs.

HLA alleles keep their responses to T-cell epitopes that are highly variable among ethnic groups. T-cell epitopes are linked with multiple alleles to increase population coverage. To anticipate the worldwide distribution of the alleles, the HTL and CTL epitopes were chosen based on their HLA alleles. The findings revealed that chosen epitopes and alleles cover a wide range of geographical regions around the world. The considered epitopes represent 84.38% of the world’s population. With 93.62% of the population, Germany is the most populous country. SARS-CoV-2 epidemics occurred in large numbers in, Spain, France and Iran. Vaccine candidates are therefore critical for protecting people in these areas from SARS-CoV-2 infection. In China, where virus originally appeared and had multiple outbreaks, population coverage was 60.83 percent.

From CTL, HTL, and B cell epitopes, vaccination candidates were chosen based on their antigenicity, toxicity, immunogenicity, population coverage, and allergenicity. The MESV was created by joining the HTL, CTL, and B cell epitopes using GPGPG, AAY, and KK linkers, respectively. Linkers are introduced as a necessary component in the evolution of MESV to improve folding, stability, and expression. When used alone, multi-epitope-based vaccinations are ineffective and require adjuvant coupling. Adjuvants are components in vaccination formulations that protect against infection while also influencing immune responses, antigen development, stability, and durability (Lee & Nguyen, 2015; Lee & Nguyen, 2015; Lee & Nguyen, 2015 As a result, a 45-amino-acid long adjuvant-defensins was combined with a 5-amino-acid linker at the N-terminus of the EAAAK linker. The EAAAK linker is used to engage the first epitope and adjuvant to aid in efficient separation of the bi-functional fusion protein domains. The final vaccination stretch was revealed to be 276 amino acids long after adjuvant and linkers were added.

The MESV construct has been found to be stable, basic, and hydrophobic based on its physiochemical features. According to the theoretical pI value, MESV was basic, ensuring a consistent physiological pH interaction. According to the estimated aliphatic index and instability index scores, the vaccination protein appears to be stable and thermostable. Its hydrophobic nature is indicated by a positive grand average of hydropathy score. MESV has been discovered to be immunogenic, antigenic, and allergenic. This points to the epitopic vaccine’s capacity to stimulate a significant immune response without causing allergic responses.

The 3D structure prediction helps give a lot of information about how important protein components are arranged in space, which is useful for studying ligand interactions, protein activities, dynamics, and other proteins (Sars-cov-, 2021; Tahir Ul Qamar et al., 2019). The MESV construct’s desired properties significantly improved after modification. The Ramachandran plot analysis reveals that the majority of residues are found in the preferred and allowed regions, with only a few residues in the disallowed region, indicating that the model’s overall quality is acceptable. The RMSD number, Poor Rotamers, Clash Score, and MolProbity all indicate that the proposed MESV construct is of high quality. To detect errors in the modelled MESV build, various structure validation methods were applied. The overall structure of the modified MESV is of good quality, as evidenced by the ERRAT quality factor (77.1822 %) and z-score (−0.37).

For an immune response to be triggered, there must be a good contact between the antigen molecule and the immune receptor molecules. After then, the improved MESV construct was docked against TLR3 to see if it might trigger a rapid immunological response. In molecular docking research, stable connections between MESV and TLR3 were found, and proficient binding required less energy.

Multi-epitope vaccines including B- and T-cell epitopes should potentially activate both humoral and cellular immune responses. Our vaccination produced the most IFN-with the highest levels of IL-10 and IL-2 activity. Antibodies also defend against SARS-CoV-2 in the extracellular environment. Furthermore, because a subunit vaccination has a variety of B-cell and T-cell epitopes, the irrelevant Simpson index (D) suggests a diversified immune response.

Because of mRNA codon incompatibility, which requires codon optimization for better expression, the translation effectiveness of foreign genes within the host system differs (Pandey et al., 2018). The obtained score was 1.0, and the GC content (53.2%) was likewise within the optimal range, implying that increased expression in the E. coli K-12 system is achievable. MESV in silico cloning’s major goal was to guide genetic engineers and molecular biologists on the predicted expression level or possible cloning sites in a specific expression system, such as the E. coli K12 system.

In this study, we used a next-generation vaccine design strategy to produce a MESV construct capable of eliciting immune responses against SARS-CoV-2. We anticipate that our vaccine, will trigger both humoral and cell-mediated immune responses. The interaction & binding patterns between both the receptor and the vaccination protein remained consistent and improved. Furthermore, effective immune responses were seen in real life during immunological simulation. MESV, which was carefully built using such a technique, could thus become a valuable asset in the fight against viral diseases.

Experimental procedures are used to produce initial raw data for computational/immunoinformatics approaches. The accuracy of immunoinformatics predictions can be limited by the quality of the data and the effectiveness of the computer techniques used. To ensure the genuine potential of proposed MESV to combat COVID-19, more in vivo and in vitro research is required.

Current developments in immunological bioinformatics areas had resulted in different servers and tools that can save cost and time of traditional vaccine development. The main problem in selection of immunodominant epitopes and suitable antigen candidates remains a hurdle for researchers. Though for designing a multiple epitope vaccine the antigenic epitopes prediction of a relevant protein by immunoinformatic methods are very helpful. By using Insilco cloning we will acquire a harmless SARS-CoV-2 vaccine that could trigger immune responses, Cellular, innate, and humoral. Though, the production and manufacture of vaccine is expensive and takes more time. Immunoinformatic approaches can decrease this load. Now a day’s researchers are finding different methods for development of multiepitope subunit vaccine. With the development of computational tools epitope prediction for antibodies become more meaningful and efficient.

## Conclusion

Current developments in the immunological bioinformatics areas has resulted in different servers and tools that are cost and time efficient for the traditional vaccine development. The main limitation in the selection of immunodominant epitopes and suitable antigen candidates remains a hurdle for researchers. Though for designing a multiple epitope vaccine the antigenic epitopes prediction of a relevant protein by immunoinformatic methods are very helpful.

By using Insilco cloning we will acquire a harmless SARS-CoV2 vaccine that could trigger immune responses: Cellular, innate, and humoral. Though, the production and manufacture of vaccine is expensive and takes more time. Immunoinformatic approaches can decrease this load. Now a day’s researchers are finding different methods for development of multiepitope subunit vaccine. The advancement in the computational tools for epitope prediction for antibodies have become more efficient and meaningful.

## Data Availability Statement

The findings of this study are available within the article.

## Ethics and Consent Statement

Consent was not required

## Animal Research Statement

Not Applicable

